# Loss of Bone Marrow β1/β2-Adrenergic Receptors Reprograms Host-Microbiota Interactions and Protects Against Diet-Induced Obesity

**DOI:** 10.64898/2026.03.05.707516

**Authors:** Adriana Alviter-Plata, Niousha Ahmari, Jennifer Gadient, Elizabeth Brammer-Robbins, Christopher J. Martyniuk, Jasenka Zubcevic

## Abstract

The gut ecosystem is shaped by multiple factors with the immune system being one of the major determinants in shaping its composition in health and disease. On the other hand, the immune system regulates its responses through the action of the sympathetic nervous system (SNS) in part through beta-adrenergic receptors 1/2 (ADRB1/2). In the past years, a clear link has been established between the immune system, SNS, and the modification of nutrient absorption by the gut microbiota in the development of diet-induced obesity. We have previously shown in male mice transplanted with bone marrow cells ADRB1/2 knock-out mice (KD) showed mild immunosuppression and microbiota changes. Post-recovery, mice were challenged with high-fat diet (HFD) for two weeks *ad libitum*. Our findings show that KD mice are protected against diet-induced adiposity and weight gain. Additionally, these mice showed an increase in residual calorific values and a decreased expression of the fatty acid transporter FAT/CD36. Suggesting a decreased absorption of lipids in the KD mice. Gut microbiota analysis showed that KD microbiota composition on a HFD remained stable with a significant enrichment in the *Bacteroidetes phylum*, which is depleted in obesity. This was associated with a switch from triglycerides to diglyceride fecal profile. Moreover, microbiome culture showed a decrease in triglycerides after an incubation with 0.1% of HFD lipid extract. Suggesting a potential role of the *Bacteroidetes phylum* in the metabolism of these lipids. Our findings demonstrate not only that the gut microbiota can modify nutrient absorption and susceptibility to diet-induced obesity but also that the immune system contributes to selective depletion of microbial members that would otherwise thrive on dietary lipids. Revealing a novel mechanism by which host immunity sculpts the gut ecosystem in ways that influence metabolic outcomes.

## Introduction

The gut microbiome is shaped by numerous internal and external factors, with the immune system acting as a central determinant of its composition under both healthy and dysbiotic conditions [1]. In turn, the microbiota trains and modulates immune activity [2, 3], establishing a bidirectional relationship that also involves the sympathetic nervous system (SNS). Through adrenergic receptor signaling, the SNS regulates immune responses [4, 5] and, when overactivated - as frequently observed in obesity [6–8]-can further disrupt microbial communities [9, 10]. Obesity is additionally characterized by chronic low-grade inflammation, which contributes to gut dysbiosis and loss of protective phyla such as *Bacteroidetes* [11–13]. Beyond immunity, the microbiota metabolizes dietary nutrients [14–16], influences nutrient absorption by altering transporter expression [17, 18], and modulates host enzymes such as lipases critical for lipid digestion [19] . In diet-induced obesity, upregulation of long-chain fatty acid transporters like FAT/CD36 has been observed [20, 21], implicating its role in the phenotype. Despite these insights, the integrated roles of the immune system, SNS, and gut microbiota in obesity remain incompletely understood.

Our previous work with a unique bone marrow (BM) chimera mouse model demonstrated that loss of adrenergic receptors in BM-derived hematopoietic cells dampened immune responses, including reduced intestinal immune cell infiltration, and produced a striking shift in the gut microbiota, highlighted by an expansion of the *Bacteroidetes* phylum [22] and decreased intestinal expression of intestinal FAT/CD36 [22], a fatty acid transporter, suggesting protection against diet-induced obesity. To test this, we challenged the chimera and their experimental controls with a two-week administration of high-fat diet (HFD), hypothesizing that its attenuated inflammatory profile would mitigate HFD-driven effects. Indeed, the BM chimera was significantly protected from HFD-induced weight gain, visceral adiposity, and blood pressure increase. These benefits were associated with resistance to HFD-induced dysbiosis at both compositional and functional levels, reduced FAT/CD36 expression relative to control chimeras on HFD, and fecal lipidomic shifts consistent with enhanced bacterial fatty acid metabolism.

Together, these findings underscore the complex, interdependent relationship between the immune system, SNS, and gut microbiota in obesity. These data further suggest that targeted modulation of the gut microbiome may provide protection against the detrimental effects of chronic sympathetic overactivation and immune dysregulation.

## Materials and Methods

### Animal Care

All experimental procedures were approved by the University of South Florida and the University of Florida Institute of Animal Care and Use Committees and complied with the standards stated in the National Institutes of Health Guide for the Care and Use of Laboratory Animals. All mice were housed in a temperature-controlled room (22–23°C) on a 12:12 h light-dark cycle, in specific pathogen-free cages and had access to standard chow mouse food and water *ad libitum* throughout the experiment. Following all *in vivo* experimental procedures mice were euthanized by isoflurane overdose followed by decapitation.

### Generation and Characterization of Bone Marrow Chimera Mice

Bone marrow (BM) chimera mice were generated as previously described [22]. Briefly, 12-week-old male ADRB/ADRB2 knock out (KO) mice (n=4), null for both the ADRβ1 and ADRβ2 genes (Jackson Laboratory, stock number 003810), were bred to 12-week-old female C57BL/6J (Jackson Laboratory, stock number 000664, n=4) to generate heterozygous ADRB1/ADRB2 KO mice. At 8 weeks old, male ADRB1/ADRB2 KO mice (n=17) were euthanized. Femurs and tibias were removed, cleaned with sterile gaze, cut, and the BM was flushed with a sterile phosphate-buffered saline (PBS) solution (Thermo Fisher, cat# 10010023). BM was pooled, homogenized gently using a pipette, and strained though a nylon cell strainer (Thermo Fisher, cat # 22363547) to leave a single-cell suspension. BM suspension was centrifuged and resuspended in PBS to adjust density of cells to 5 × 10^6^ BM cells/ml of PBS. 200µl of BM cells in suspension from ADRB1/ABDB2 KO mice were reconstituted to x-ray-irradiated (950 Rad) 6-week-old male C57BL/6J wild type (WT) mice by a single tail vein injection at a ratio of 1:4 (donor to recipient), to generate the BM ADRB1/ADRB2 knock-down Chimera mice (KD Chimera, n=10). At the same time, control WT chimera mice (WT Chimera) were generated by reconstitution of C57BL/6J whole BM cells into x-ray-irradiated age- and sex-matched C57BL/6J mice using the same protocol (n= 13). All reconstituted mice were maintained on antibiotic drinking water (Baytril 0.25 mg/mL, Bayer HealthCare Animal Health Division, Shawnee Mission, KS) throughout the 3 weeks of recovery. Following this, all chimera mice were allowed an additional 9 weeks of recovery to allow full bone marrow reconstitution. Successful reconstitution was confirmed by ADRβ1/2 genotyping in circulating mononuclear cells (MNCs) as before [22]. Briefly, genomic DNA was isolated from peripheral blood MNCs following digestion at 55°C in buffer containing 50 mM Tris-HCl (pH 8.0), 10 mM EDTA, 100 mM NaCl, 0.1% SDS and 1mg/ml proteinase K. PCR was performed to check for presence of mutant (ADRβ1/2 KO) probe using the primer pairs the following primer pairs: F-*TTG GGA AGA CAA TAG CAG GC*; R-*CGC TGT CCA CAG TGG TTG T*.

### Administration of High Fat Diet in KD and WT Chimera

Following recovery and confirmation of genotype, select KD and WT Chimera mice (n=5-8 per group) were administered high-fat diet (HFD) containing 60 kcal% fat and 20 kcal% carbohydrates (D12492, Research Diets Inc., New Brunswick, NJ) for two weeks ad libitum. Separate KD and WT Chimera mice (n= 5-8 per group) were fed a control rodent diet (CNT) containing 10 kcal% fat and 20 kcal% carbohydrates (D12450J, Research Diets Inc., New Brunswick, NJ). Mice in both groups had access to water *ad libitum*.

### Body weight and Food Intake in WT and KD Chimera on Control and High-Fat Diet

Mice were group-housed based on genotype and diet (n=2-4/ cage). Food pellets were weighed once per day for each cage throughout the experiment and averaged for each week to determine the cage food intake. Body weight was measured in each mouse prior to administration of CNT and HFD, and then once per week throughout the experiment. Data are presented as means ± SEM.

### Fat Pad Collection

At endpoint, all mice were euthanized by isoflurane overdose (n= 5-8/ group) (4% in room air) followed by decapitation. For collection of brown fat pads, clean forceps were used to elevate the skin between the shoulder blades. A small incision was made in the skin to expose brown fat. Total brown fat pads were harvested and weighed. Following this, the abdominal wall was cut open from the genitals to the rib cage. Retroperitoneal fat pads surrounding the kidney to isolate and perirenal fat. The adrenal gland was removed from the retroperitoneal fat pads prior to weighing. Care was taken to collect all visible fat for accuracy as per [23].

### Blood Pressure Measurements in Mice

Blood pressure measurements were taken in all mice (n= 5-8 per group) on two consecutive days per week throughout the experimental paradigm. Mice were placed in metal magnetic restraints attached to a pre-heated platform (37 °C). After 10 minutes of acclimatization, a tail cuff with a pneumatic pulse sensor was attached to the tail (Visitech Systems, Inc.). Mice were allowed to habituate to the tail cuff measurement for 10 readings. Ten to fifteen additional readings were performed every 90 seconds. Systolic blood pressure (SBP) values were recorded on Model BP200 (Visitech Systems, Inc.). Readings were accessed for validity, and valid readings were averaged to obtain an average SBP for each mouse. Readings were taken at 10-11am on two consecutive days per week and averaged to obtain a weekly time point for each mouse.

### Residual Calorimetric Analysis

Fecal samples were collected from all groups at endpoint (n= 5-6/ group), weighed and fully dehydrated by placing them at 60°C for 48h followed by 24h at 80°C, confirmed by weighing. For fecal calorimetric measurements, dried samples were placed on a thermogravimetric analyzer differential calorimeter (TA Instruments Q600) and flooded with an N_2_. Temperature was increased by 5° C per minute until it reached 550°C. Energy transfer was captured, and heat flow was analyzed using Thermal Advantage (v5.5.24). Linear integration peak (maximum energy transfer from the sample) was measured between the limits of 200°C and 550°C. To calculate the residual calories in each group and how diet affected this, caloric values for feces from HFD mice were normalized to caloric values of feces from CNT mice within each group, using the following equation:

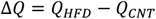

### RNA Isolation and RT-qPCR in the proximal colon

RNA isolation from the proximal colon (n= 3-6/ group x2 technical replicates) was performed as previously described [24]. Real time quantitative PCR (RT-qPCR) was used to measure relative expression levels of FAT/CD36 using the following forward and reverse primers: F- *CCA GTG TAT ATG TAG GCT CAT CCA*, R- *TGG CCT TAC TTG GGA TTG G* (IDT) [25]. Expression data were normalized for the expression of housekeeping gene GAPDH (F- *CTC TGG AAA GCT GTG GCG TGA T*, R- *CAT GCC AGT GAG CTT CCC GTT CAG*) (IDT) [26]. RT-qPCR was performed using a CFX96 Touch Real-Time PCR Detection System (Bio-Rad, Hercules, CA). Relative gene expression was assessed using the relative ΔΔCt for comparisons of more than two groups, where Ct represented the threshold cycle.

### Bacterial Sequencing and Analysis of Cecal Bacterial Communities

Cecal bacterial samples (n= 3-8/ group) were collected from all mice at end point. Bacterial DNA was extracted using ZR Fecal DNA MiniPrep (Zymo Research, Irvine, CA) as per manufacturer’s protocols. DNA concentrations were determined using the NanoDrop-2000 spectrophotometer (Thermo Scientific). Bacterial 16S rRNA sequencing was performed on the Illumina platform using Miseq v2 reagent kit (Illumina, Inc., San Diego, CA) as previously reported [27]. Sequenced reads were demultiplexed according to a combination of forward and reverse indices. Demultiplexed reads were analyzed with “Quantitative Insights into Microbial Ecology 2” (QIIME 2) [v.2023.2] [28, 29]. DADA2 plugin was used to merge paired end fastq files, denoising, chimera removal and construction of amplicon sequence variants (ASV). Taxonomy was analyzed using the Silva v.132 databases at 99% identity pre-trained for the V3-V4 region [30]. Rarefaction was performed at a sampling depth of 14529 sequences per sample. Alpha-diversity was estimated using Shannon metric. Linear discriminant analysis (LDA) effect size (LEfSe) was performed using the R package MicrobiomeMaker (v. 1.10) with a false discovery rate adjusted to 0.01 and an LDA score equal to or above 4. The software PICRUSt2 (version 2.3.0b) was used with default settings to infer approximate functional potential of the microbial communities in all groups. Statistical analysis of functional profiles was done using STAMP [31] (v.2.13). with Benjamin-Hochberg FDR was used for multiple test correction.

### Short Chain Fatty Acid (SCFA) Extraction and High-Performance Liquid Chromatography (HPLC)

Short chain fatty acids (SCFAs) were extracted from cecal samples (n= 4-8/ group) as per our published protocols [24]. Briefly, 200 μL of HCl was added to the cecal samples to preserve the volatile SCFAs, and vortexed vigorously for even suspension. Five milliliters of methylene chloride solution was added, and samples were gently rotated at RT for 20 min. After centrifugation at 3,500 rpm for 5 min, the supernatant was discarded. Five hundred microliters of 1N NaOH was added to the organic pellet, followed by an additional 20 min rotation at room temperature. After spinning at 3500 rpm for 5 min, the top aqueous phase was collected and mixed with 100 μL of HCl before being filtered for injection into the HPLC column. SCFA concentrations were calculated by using the area ratios obtained from the biological samples, average signal of the blank samples as intercept, and the slopes obtained from the analysis of the calibration series samples.

### Lipid Extraction, Identification and Lipidomic Analysis

Lipidomic analyses were performed in fecal and serum samples at endpoint and in food pellets from CNT and HFD [32–34] for comparison. Briefly, samples were lyophilized, and lipids were extracted using the Bligh and Dyer method [35]. Lipid concentrates were run through a liquid chromatography system (LC) coupled with mass spectrometry (MS) for separation and identification.

Samples were analyzed with MetaboAnalyst (v 6.0) as previously reported in [28]. Briefly, samples were normalized using Quantile normalization on relative lipid abundance in food samples or dry weight for fecal lipids. This was followed by Log transformation (base 10). Auto scaling (mean-centered / divided by the standard deviation of each variable) was then conducted to normalize data. A Volcano Plot was generated considering a fold change value of 2.0 (raw p-value <0.05).

### Whole Microbiome Culture

End point fecal samples were collected from WT and BM chimera on HFD and used for whole microbiome cultures in Gifu Anaerobic Media (GAM) as described in [36]. Briefly, fecal pellets from two groups separately were resuspended in 1X sterile phosphate buffered saline (PBS). Debris was allowed to settle in the bottom and 50 μL of bacteria were inoculated in 40 mL of sterile deoxygenated GAM broth. Bacteria were cultured for 24h following which an aliquot from each group was reintroduced to fresh GAM media with 0.1% HFD lipid extract for another 24h. Bacterial metabolites were collected and filtered through a 0.22 μm membrane, which were further used for lipidomic analysis as described previously.

### Lipase activity Assay

Colons were collected at end point (n= 3-6/ group x2 technical replicates) for the analysis of lipase activity (Abcam ab102524) following the manufacturer instructions. Briefly, frozen colons were weighed, homogenized in liquid nitrogen (N_2_) and resuspended in assay buffer. Samples were centrifuged and supernatant was collected for the lipase measurements. Kinetic synthesis of glycerol was determined by measurements at 570 nm every 2 min for 90 minutes at 37°C, and the sample enzymatic activity was calculated as amount of glycerol synthesized within the linear part of the enzyme activity curve. Lastly, average enzyme activity was normalized to tissue weight.

### Statistical Analysis

After testing for normality (Shapiro–Wilk test), intergroup differences were analyzed by unpaired two-tailed Student’s t-test for single comparison, or by one-way or two-way analysis of variance (ANOVA) with an appropriate multiple-comparison test, as indicated in the figure legends. The F test for equality of variances was performed and, if indicated, appropriate corrections were made (Welch’s correction for the t-test) using Prism 9 (GraphPad). Data are presented as mean ± SEM. For microbiota analysis, linear discriminant analysis (LDA) effect size (LEfSe) was performed using the R package MicrobiomeMaker (v. 1.10) with a false discovery rate adjusted to 0.01 and an LDA score equal to or above 4. Statistical analysis of functional profiles was done using STAMP [31] (v.2.13). with Benjamin-Hochberg FDR was used for multiple test correction.

For lipidomics analysis samples were analyzed as reported previously [28] with MetaboAnalyst (v 6.0) [37]. Lipid data were first normalized using Quantile normalization on relative lipid abundance in each sample, followed by Log transformation (base 10). Auto scaling (mean-centered / divided by the standard deviation of each variable) was then conducted to normalize data. A Volcano Plot was generated considering a fold change value of 2.0 (raw p-value <0.05).

## Results

### KD chimera mice are protected against dietary fat-induced weight gain, visceral fat accumulation and rise in blood pressure

WT and KD chimera mice were administered either CNT or HFD for two weeks (Fig. 1A) to determine the dietary effects on adipose expansion, weight gain, and blood pressure. While there was no significant difference in total food intake between the groups (Fig. 1B), the WT on a HFD showed a significant body weight increase from baseline compared to WT on CNT diet, which was not observed in the KD mice (Fig. 1C). Moreover, HFD caused a significant expansion in visceral fat pad weights (epididymal and retroperitoneal fat pads) but only in WT mice on HFD, with no effects seen in normal adipose tissue (Fig. 1D-1F). Additionally, systolic blood pressure (SBP) was higher in the WT compared to KD chimera following two weeks of HFD intake (Fig. 1G), suggesting overall protection against effects of HFD in the KD chimera.

**Figure 1.**
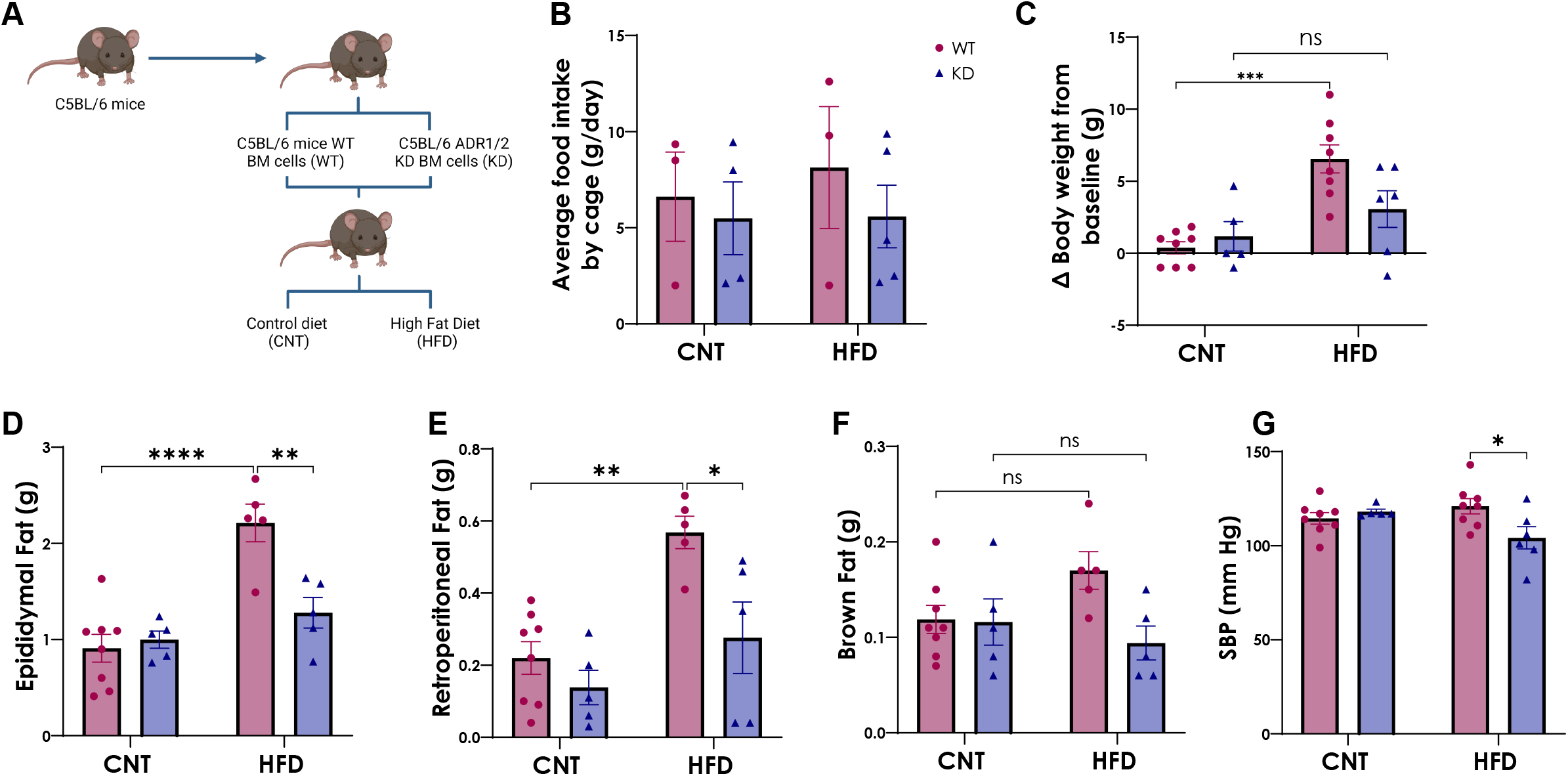
Murinometric parameters of WT and KD on CNT and HFD. **A**. Schematic representation of bone marrow transplant on C5BL/6 mice. C5BL/6 mice received whole bone marrow cells from same strain mice to generate wild-type group (WT) or ADRB1/ADRB2 KO mice to generate knock-down group (KD). These groups were further divided and were either administered for two weeks with either a high-fat diet (HFD), to generate WT on HFD or KD on HFD, or a control diet (CNT), to generate WT on CNT or KD on CNT. **B**. Average food intake by cage (g/day). **C**. Difference in body weight from baseline (g). **D**. Brown fat weight collected at endpoint (g). **E** and **F**. Fat pad collected from retroperitoneal region and retroperitoneal region prior removal of adrenal glands (g). **G**. Systolic blood pressure measured by non-invasive tail cuff (mmHg). Data is represented as Mean ± SEM, with Tuckey post-hoc *p< 0.05, **p< 0.01, ***p<0.001, 2way ANOVA. **Abbreviations: WT-** Wild-type; **KD-** Knock-down; **BM-** Bone marrow; **ADR1/2-** Adrenergic β receptor 1 and 2; **CNT-** Control; **HFD-** High-fat diet; **SBP-** Systolic blood pressure.

### Higher fecal calorific values are associated with reduced relative expression of colonic FAT/CD36 in the KD chimera on HFD

Fecal calorimetric analysis was performed at endpoint to determine residual calorific values suggestive of potential deregulation in absorption of dietary fatty acids (Fig. 2A-2D). We observed higher residual calorific values in the KD compared to WT mice (Fig. 2E). Moreover, FAT/CD36 colonic transcript was higher in the WT compared to the KD mice following HFD (Fig 2F), suggesting reduced absorption of lipids in the KD chimera despite equivalent food intake between the groups (Fig1B).

**Figure 2.**
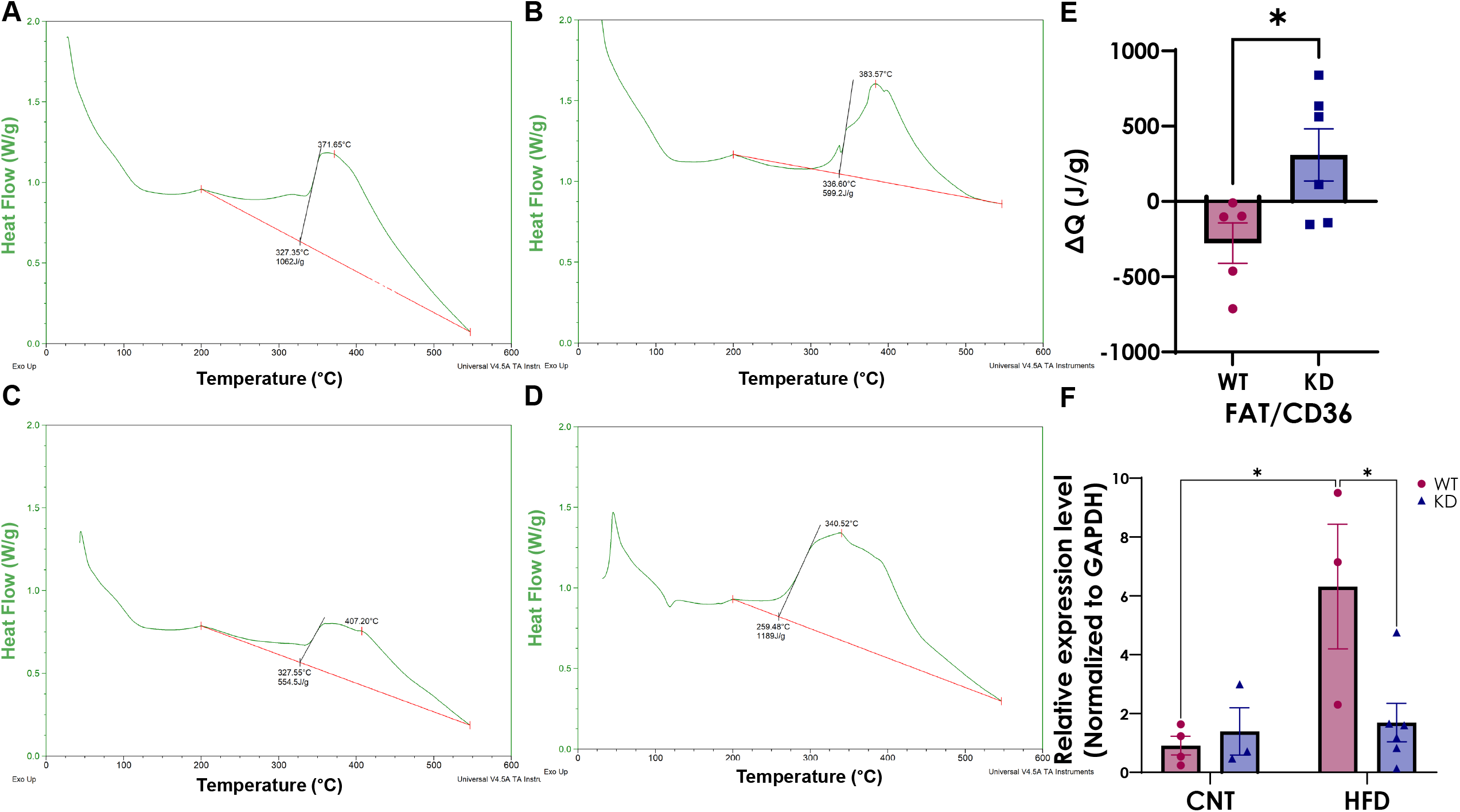
Fecal energetic profile graphs normalized to weight in **A**. WT on CNT, **B**. WT on HFD, **C**. KD on CNT, and **D**. KD on HFD. **E**. Residual fecal calorific values normalized to CNT diet in WT and KD groups. **F**. Relative expression of colonic FAT/CD36 normalized to expression of GAPDH. Data is represented as Mean ± SEM (N=5-6/group; *p< 0.05, **p< 0.01, ***p<0.001, Ttest). **Abbreviations: WT-** Wild-type; **KD-** Knock-down; **CNT-** Control; **HFD-** High-fat diet; **FAT/CD36-**Fatty acid translocase/ cluster of differentiation 36

### HFD produced an enrichment in *Bacteriodales* in the KD but not WT mice but no difference in SCFAs

Our previous work showed marked differences in the gut bacterial composition and abundances in the KD model [22], marked by an enrichment in the *Bacteroidetes phylum*. To investigate how HFD may affect this, we sequenced the cecal content in all mice at endpoint. In contrast to the WT, HFD challenge did not alter the overall diversity and richness in the KD chimera as measured by the alpha diversity index (Fig. 3A). Coinciding with our previous findings we observed a significant enrichment of the *Bacteroidales* order and its Bacteroidetes *phylum* in KD mice compared to WT on HFD (Fig. 3B), suggesting a degree of stability of the KD bacteria to HFD. To investigate potential functional differences, we performed predictive metagenomics analysis with PICRUSt [38]. We observed overall fewer changes in the gut microbiome function in the KD mice compared to WT on HFD (Fig. 3C-3D, Supplementary Table 1). In addition, we observed an elevation in the relative frequency of fatty acid elongation pathway (FASYN. ELONG. PWY) in the KD mice following HFD (Fig 3E), often linked to bacterial phospholipid synthesis [39]. Lastly, no difference was observed in fecal concentrations of the main short-chain fatty acids (acetic, butyric, and propionic acid) between the two groups (Fig 3F-3H).

**Figure 3.**
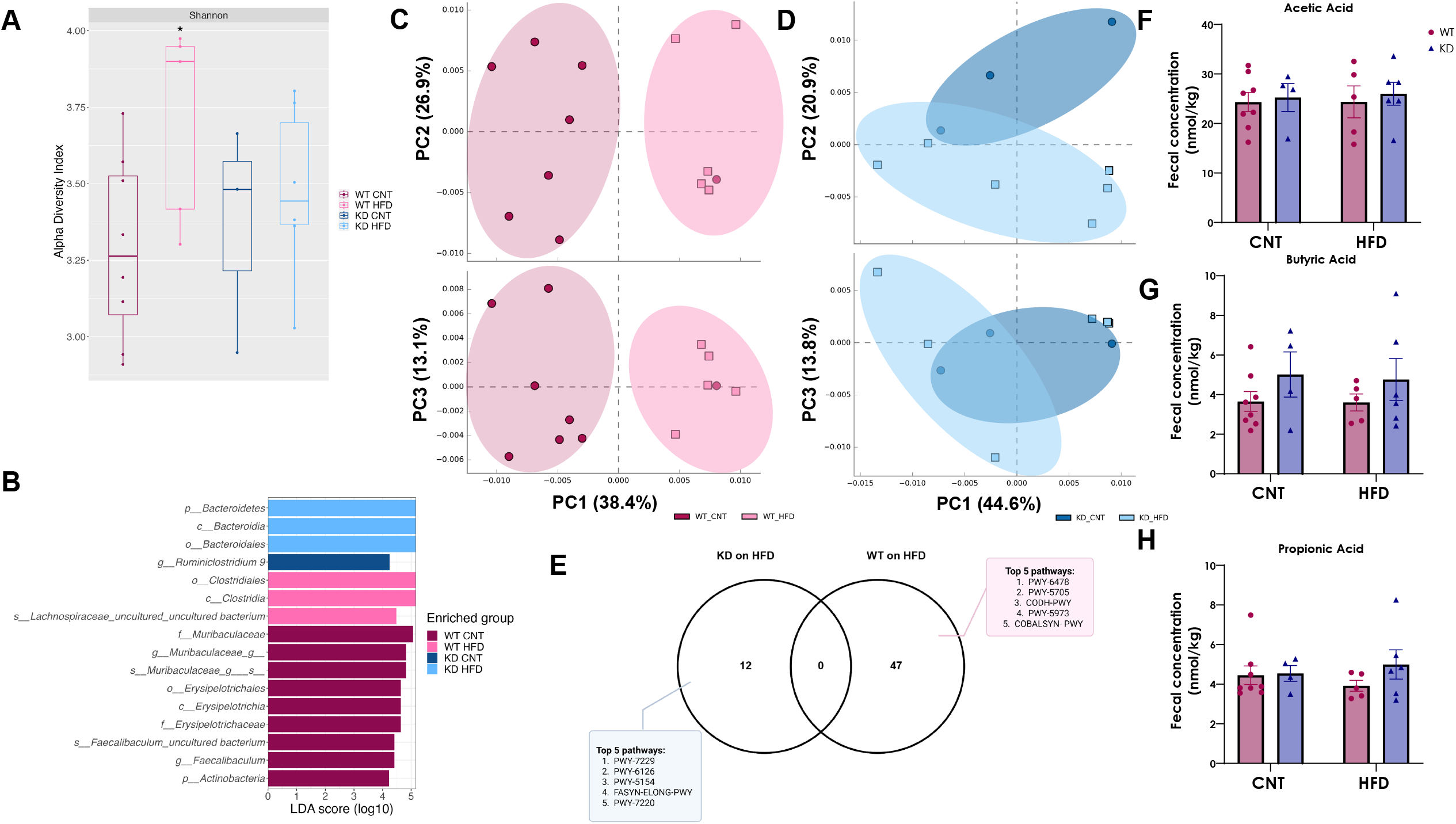
Colonic microbial communities’ analysis in WT and KD groups. **A**. Alpha-diversity boxplot represented by measured of Shannon index. *p < 0.05, 1way ANOVA vs WT CNT. **B** Linear discriminant analysis effect size (LEfSe) with false discovery rate–adjusted P<0.01 and linear discriminant analysis (LDA) ≥4.0. Principal Component Analysis (PCoA) Plots of predictive metagenomic functions of **C**. WT on CNT and HFD and **D**. KD on CNT and HFD. **E**. Venn Diagram of significantly higher predictive pathways in HFD in both WT and KD. Fecal concentration of main SCFAs, **F**. acetic acid, **G**. butyric acid and **H**. propionic acid. Data is represented as Mean ± SEM, with Tuckey post-hoc *p< 0.05, **p< 0.01, ***p<0.001, 2way ANOVA. **Abbreviations: WT-** Wild-type; **KD-** Knock-down; **CNT-** Control; **HFD-** High-fat diet; **SCFA-** Short-chain fatty acid.

### Shifts in the lipidomic profiles in the feces of KD mice on HFD suggest elevated bacterial metabolism of dietary triglycerides

To investigate if KD gut bacteria may utilize dietary lipids, we performed shotgun lipidomics in fecal and serum samples of all mice at endpoint and compared to the lipid profiles in the HFD pellets. We observed a difference in abundances of dietary triglycerides (TAG) and diglycerides (DAG) in the WT compared to KD mice on HFD (Fig 4A-4F, Supplementary Table 2) despite equivalent food intake (Fig 1C). To discard any potential difference in the overall relative frequency of lipid classes in the diets, we also performed lipidomic analysis in food pellets. No differences were observed in the relative frequency of the different lipid classes between the two diets (Fig. 4E). A higher relative frequency of DAGs and a lower relative frequency of TAGs was observed in the KD mice on a HFD compared to the WT on the same diet (Fig 4F) with similar results in circulation (Fig. 4C and 4D). Moreover, a higher relative frequency of phosphtatidylinositols (PI) and phosphatidylserine (PS) were observed in the KD compared to WT on HFD (Fig 4D), suggesting elevated utilization of TAGs by the KD gut bacteria. To test this directly, equal weights of endpoint fecal samples from WT and KD mice were cultured in the presence of 0.1% of HFD lipid extract overnight and performed shotgun lipidomics in the supernatant. Analyses showed a lower TAG to DAG proportion in the supernatant of KD compared to WT whole bacteria, suggesting elevated bacterial utilization of dietary TAGs in the KD mice (Fig 4G and 4H).

**Figure 4.**
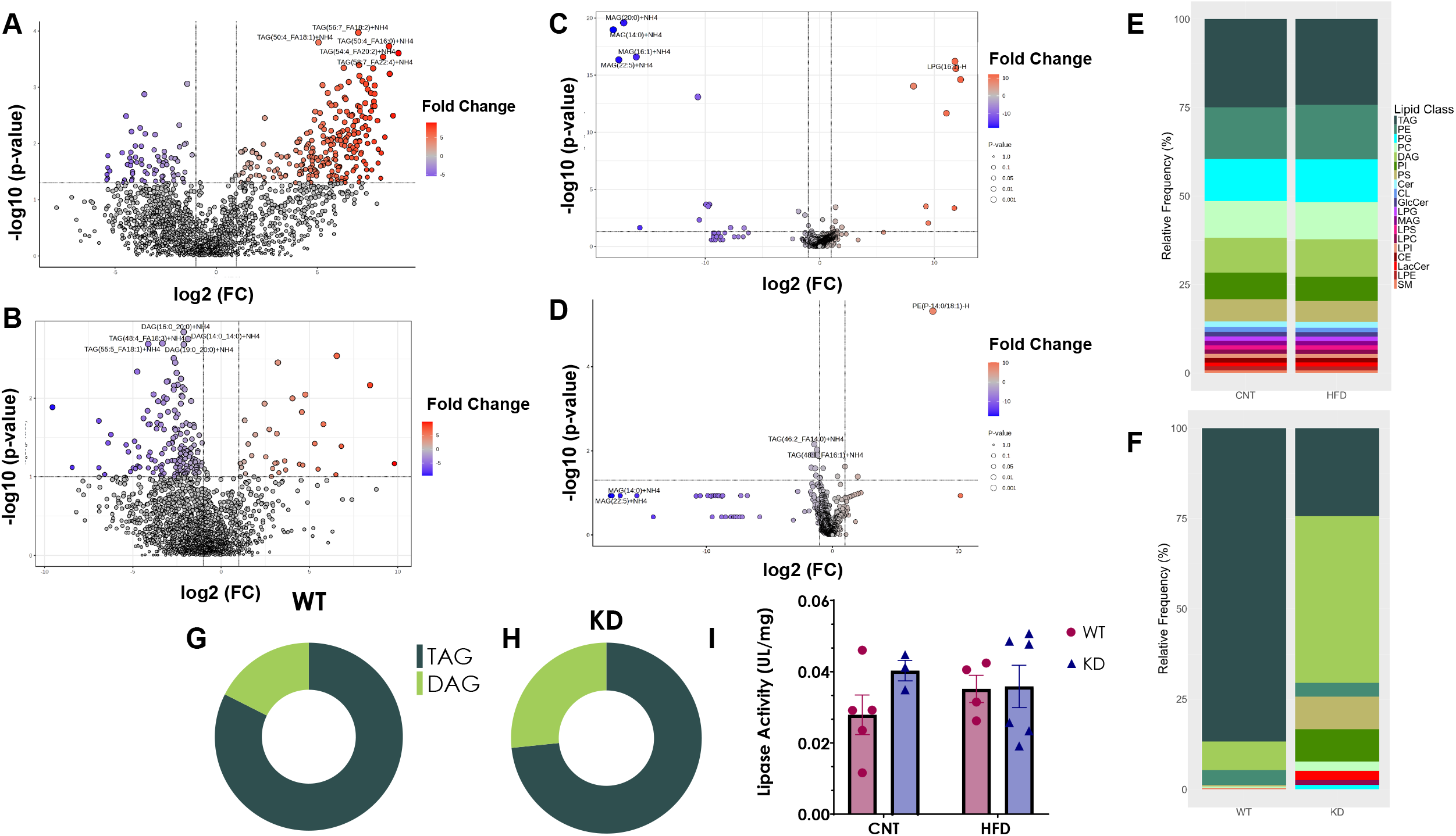
Fecal lipidomic analysis of WT and KD on a HFD challenge. **A**. Volcano plots of A. WT on HFD and **B**. KD on HFD of the fecal the differentially abundant lipids (p < 0.05) outlined in red and blue. Volcano plots of **C**. WT on HFD and **D**. KD on HFD of the differentially abundant lipids (p < 0.05) outlined in red and blue in serum **E**. Lipid distribution represented as relative frequency of lipid classes present in HFD and CNT diets. **F**. Lipid distribution represented as relative frequency of differentially abundant lipids in HFD in both WT and KD fecal samples. **G**. and **H**. TAG and DAG proportions in whole microbiome cultures from WT and KD on CNT incubated with 0.1% of lipids extracted from HFD respectively. **I**. Colonic lipases activity normalized to tissue weight. Data is represented as Mean ± SEM, with Tuckey post-hoc *p< 0.05, **p< 0.01, ***p<0.001, 2way ANOVA. **Abbreviations: WT-** Wild-type; **KD-** Knock-down; **CNT-** Control; **HFD-** High-fat diet; **TAG-**Triacylglycerols; **PE-** Phosphatidylethanolamines; **PG-** Phosphtatidylglycerols; **PC-** Phosphatidylcholines; **DAG-** Diacylglycerols; **PI-** Phosphtatidylinositols; **PS-** Phosphatidylserine; **Cer-** Ceramides; **CL-** Cardiolipins; **GlcCer-** Glucosyl ceramides; **LPG-** Lyso Phosphtatidylglycerols; **MAG-** Monoacylglycerols; **LPS-** Lyso Phosphatidylserine; **LPC-** Lyso Phosphatidylcholines; **LPI-** Lyso Phosphtatidylinositols; **CE-** Cholesteryl esters; **LacCer-** Lactosyl ceramides; **LPE-** Lyso Phosphatidylethanolamines; **SM-** Sphingomyelins.

### HFD produces no changes in the activity of lipases activity

To investigate any influence of the host in the overall fecal lipidomic profile we performed a colonic lipase activity assay. No difference in colonic lipase activity was observed between the two groups on HFD (Fig 4I), thereby excluding the effects of host lipases on the observed fecal lipids profiles, thus highlighting the influence of bacteria.

## Discussion

C57BL/6J mice, which have the same genetic background as our chimeric mice, are considered a model of diet-induced obesity. Temporal studies on the effect of the administration of HFD on this mouse strain have shown that HFD causes a rapid increase in their body weight starting at week 2 [40]. Contrasting to what has been reported, the KD chimera mouse, characterized by reduced immune responses [22], did not have a significant increase in body weight and expansion of adipose tissue. Overall, this suggests that the KD mice are somewhat protected from both weight gain and expansion of visceral fat despite same food intake and colonic lipase activity. In the context of obesity, HFD can significantly increase the production of pro-inflammatory cytokines like TNF-α in the ileum in the time frames investigated here [41]. Additionally, HFD leads to a chronic low state of inflammation in adipose tissue, often characterized by a polarization of macrophages towards a pro-inflammatory profile [42]. Although we observed no differences in the expression of inflammatory cytokines in our model, the chimera mice are characterized by a reduced presence of activated macrophages at baseline [22], with the caveat that we did not measure the effects of HFD on circulating macrophages in the current study.

We previously demonstrated that reduced immune system activity due to decreased BM adrenergic receptor signaling had a significant effect on the gut ecosystem in the KD chimera [22]. The observed shifts at baseline extended to administration of HFD as measured by analyses of bacterial diversity. These effects were further associated with a reduced expression of FAT/CD36 and increased fecal calorific values, independent of food intake or host lipases levels, suggesting reduced absorption of dietary lipids by the KD chimera. Current evidence shows that expression of the FAT/CD36 transporter is upregulated in diet-induced obesity [20]. While there is no information regarding the effect of microbiota on colonic FAT/CD36, reports link gut microbiota to changes in expression of this transporter in skeletal tissue and liver [17, 18]. Furthermore, the expression of FAT/CD36 can be affected by inflammatory markers that in turn can modify the gut microbiota composition directly or indirectly [43, 44]. However, precise mechanisms of microbiota-host interactions that contribute to regulation of FAT/CD36 expression are unknown and warrant further investigations.

Additional to a protection to diet-induced weight gain our KD mice were protected against the deleterious effects of HFD on blood pressure. Hyperactivation of SNS can lead to altered immune responses in hypertension [4], and this effect can precede the increase in blood pressure [5]. Decreased SNS-immune communication in our KD model may in part be contributing to this effect. KD mice on a HFD had a higher relative frequency of the *Bacteroidetes phylum*, which is depleted in hypertension [27] and can reportedly protect against increase in blood pressure[27], although precise mechanisms are unclear. A significant expansion in the *Bacteroidetes phylum* in our KD chimera [22] was sustained after a HFD challenge in the current study. In contrast, this *phylum* is depleted in obesity and following a HFD in humans and WT mice [11–13] similarly to what we observed in the current study, highlighting the importance of this *phylum* in the maintenance of a lean phenotype. While several factors can contribute to the depletion of *Bacteroides* in obesity, the immune system plays a role in the development of adverse conditions in the gut that can contribute to diet-induced dysbiosis via elimination of select gut bacteria including members of the *Bacteroidetes phylum* [45]. Additionally, some evidence shows an interaction between this *phylum* and the expression of FAT/C36 [46, 47]. However, this is species-specific, and our future studies will investigate direct modulation of FAT/CD36 by select *Bacteroidetes* species.

The expansion in *Bacteriodetes* was associated with an increase in the frequency of FASYN.ELON.PWY, a functional bacterial pathway that predicts elongation of the lipid carbon chain in the synthesis of bacterial membrane [39, 48]. As we observe a decreased TAG to DAG ratios in fecal samples from the KD chimera on HFD *in vivo*, and following culturing of whole KD microbiota with lipid extracts *in vitro*, our results suggest an association between the abundances in *Bacteroidetes phylum* and bacterial utilization of dietary TAGs that warrants exploration in future studies.

Our findings underscore the critical role of the neuro–immune–gut axis in the development of diet-induced obesity. Nonetheless, some limitations should be noted. First, our study was restricted to male BM chimera mice. Although no weight-gain in obesity have similar prevalence in males and females [49], fat distribution accumulation changes between male and female mice [50], which may influence outcomes. Additionally, microbiota composition is sex specific, highlighting the need to address sex differences in future studies. Our analyses also focused on nutrient excretion rather than direct absorption, and future studies will assess the role of gut bacteria on dietary nutrient absorption.

## Conclusion

Our findings advance the field by uncovering a previously underappreciated dimension of host– microbiome interactions. We demonstrate not only that the gut microbiota can shape nutrient absorption and susceptibility to diet-induced obesity, but also that the immune system actively contributes to the selective depletion of microbial members that would otherwise thrive on dietary lipids. This reveals a novel mechanism by which host immunity sculpts the gut ecosystem in ways that influence metabolic outcomes. Importantly, our work highlights the intricate cross-talk between the immune system, the sympathetic nervous system, and the gut microbiome in driving obesity pathogenesis. These insights point to microbiota modulation as a powerful and innovative therapeutic strategy to buffer against the harmful consequences of immune and sympathetic overactivation, with broad implications for the treatment and prevention of obesity and related metabolic disorders.

## Data Availability

Sequencing data is available at PRJNA1354833 https://www.ncbi.nlm.nih.gov/bioproject/PRJNA1354833.

## Acknowledgements

We wish to acknowledge Dr. Kristin Kirschbaum, Director of the Instrumentation Center at the University of Toledo, who facilitated performance of residual calorimetric analysis experiment. We thank the ATCL for its LC-MS/MS data analysis, whose funding was obtained from NIH Shared Instrumentation Grant #S10OD036420

## Author Contributions

A.A.P. and N.A. performed all the experiments. E.B.R. and C.J.M. performed lipidomics and helped with lipidomics analyses and interpretations. J.G. trained A.A.P. in performing residual fecal calorific analyses, provided the instrumentation and helped with interpretation of results. A.A.P. performed all the bioinformatics analysis under supervision of C.J.M. and J.Z. A.A.P. and J.Z. drafted the manuscript; A.A.P., C.J.M. and J.Z. edited and revised manuscript; J.Z. funded the study. All authors approved the final version of manuscript.

